# Large time step discrete-time modeling of sharp wave activity in hippocampal area CA3

**DOI:** 10.1101/303917

**Authors:** Paola Malerba, Nikolai F. Rulkov, Maxim Bazhenov

## Abstract

Reduced models of neuronal spiking activity simulated with a fixed integration time step are frequently used in studies of spatio-temporal dynamics of neurobiological networks. The choice of fixed time step integration provides computational simplicity and efficiency, especially in cases dealing with large number of neurons and synapses operating at a different level of activity across the population at any given time. A network model tuned to generate a particular type of oscillations or wave patterns is sensitive to the intrinsic properties of neurons and synapses and, therefore, commonly susceptible to changes in the time step of integration. In this study, we analyzed a model of sharp-wave activity in the network of hippocampal area CA3, to examine how an increase of the integration time step affects network behavior and to propose adjustments of intrinsic properties of neurons and synapses that help minimize or remove the damage caused by the time step increase.

**Highlights:**

- Spiking models of neural network activity are sensitive to the integration step
- Larger integration time steps are preferable in simulating large networks
- Case study of CA3 sharp waves shows time step increase damages network dynamics
- Neuronal and synaptic parameters adjustments rescue the dynamics at large time step^1^

## 1. Introduction

Sleep is known to be beneficial for memories, in particular as a stage in which the brain is not processing input but can freely elaborate on recently learned memories, in a process called memory consolidation [1]. Oscillatory activity found in different brain regions during sleep has been shown to be influential in the process of memory consolidation and the presence and coordination of rhythms with different frequencies and time scales is hypothesized to be necessary for memory consolidation to occur [2–9]. Hence, studies of oscillations observed in the brain at various stages of sleep are very important for understanding of mechanisms of memory formation and consolidation. One of the essential rhythms within sleep oscillations is given by sharp-wave ripple (SWR) complexes. These are short-lived (with duration of 50-100ms) events observed in hippocampal activity during stages of quiet wake and slow wave sleep. These events are detected in the traces in the local field potential (LFP) measured across layers in hippocampal area CA1, where they appear as a strong deflection in stratum radiatum followed by fast (>150Hz) oscillations in stratum pyramidale. The strong deflection is caused by inputs from hippocampal are CA3 that projects its activity onto the pyramidal cells of CA1. Therefore, sharp waves are formed by the overall activity of CA3 pyramidal cells and evoke ripples through the activation of CA1 inhibitory neurons, generating fast transient local oscillations.

Understanding of dynamical mechanisms behind the onset of these type of oscillations and transitions causing qualitative changes of oscillation regimes rely on numerical simulations of large-scale network models that have to be examined in the in the multi-dimensional space of the network parameters. This parameter space usually includes parameters controlling the intrinsic dynamics of neurons, synapses and background noise. Use of reduced phenomenological models of neurons significantly reduces the complexity of such analysis by reducing the complexity of the model, the computational time and the number of control parameters. However, even with reduced network models of spiking neurons, simulations consume significant computational resources. Reduction of network simulation time could be extremely beneficial for this type of studies. A way to speed up the simulations is to write model in the form of map, for example using fixed step Euler method, and increase the integration time step while paying attention to the accuracy of the dynamical behavior of the model. The map can also be modified to control the dynamics and make corrections of errors introduced by larger integration steps. Importantly, the increase of integration time step has limits, as time resolution should be sufficient to capture the dynamics of the fastest processes in the network.

In this work, we used a recently developed model of SWR activity in CA3 and CA1 [10–13], and focused on sharp wave activity in CA3. The model included 2800 neurons aggregating the two regions, and the model equations were solved using a standard 1-step Euler integration algorithm. The complex dynamics of SWRs in the biological hippocampus as well as the co-existence of multiple spatial scales (i.e., fine resolution necessary to describe a single SWR event and large-scale network dynamics controlling multiple co-existing SWRs), require scaling computational networks to a size similar to the actual biological size of a human hippocampus (about 16 million pyramidal cells have been estimated in the CA1 area alone [14]). Therefore, new strategies are necessary to accelerate the computer simulations of network activity in this CA3-CA1 network.

We found highly nonlinear effects induced into the network activity by a large increase in the integration time step, from 0.001 to 0.5ms. We looked for the range of integration step sizes that maintained consistency of the network activity features. We further manipulated the network parameters to achieve qualitatively similar patterns of activity in the network solved with very large step size and a fine step size. In particular, we focused on preserving the emergence of the sharp waves in CA3 activity, and in maintaining their exponentially distributed in-between times.

## 2. Network Model Structure

### 2.1. Cells: equations and parameters

We model sharp wave activity in the hippocampal area CA3 using a network of 240 basket cells and 1200 pyramidal cells. Each cell type is modeled as one Adaptive-Exponential Integrate and Fire neuron [15]. All cells in the same population have the same intrinsic parameters, but different inputs (*I(t)*), which introduce heterogeneity within both excitatory and inhibitory neuron populations. The inputs are separated in a deterministic component, a noise component and a synaptic component. In the deterministic part of the input, every cell receives a different direct current (*I_DC_*) term, fixed for the duration of the simulation, with values selected from a Gaussian distribution (with parameters different for the two cell types). In the noise component of the input, every cell receives an independent Ornstein-Uhlenbeck process [16] (OU-process, *η_t_*), with cutoff at 100Hz, which roughly resembles a single-pole filtered white noise. This noisy input is added to put the single cells in a state of noise-driven spiking, which is considered a good representation of *in vivo* cell activity [17–20]. The standard deviation of the OU process 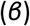 controls the size of the noisy fluctuations in the sub-threshold voltage of the cells, and is fixed within each cell type. Parameters of the deterministic and noisy input component are chosen to have cells in a noise-driven spiking regime, rather than a deterministic spiking regime perturbed by noise. For each neuron, the equations are

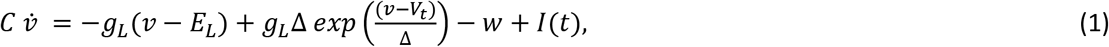

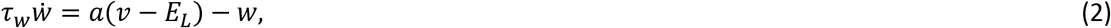

with spike reset conditions at the threshold

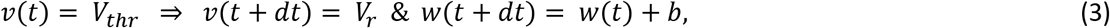

and input current that include DC bias, noise and synaptic inputs.

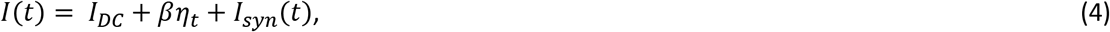

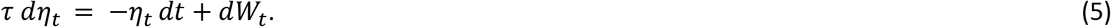

The intrinsic parameters of the neuron models are presented in Table 1.

**Table 1:**
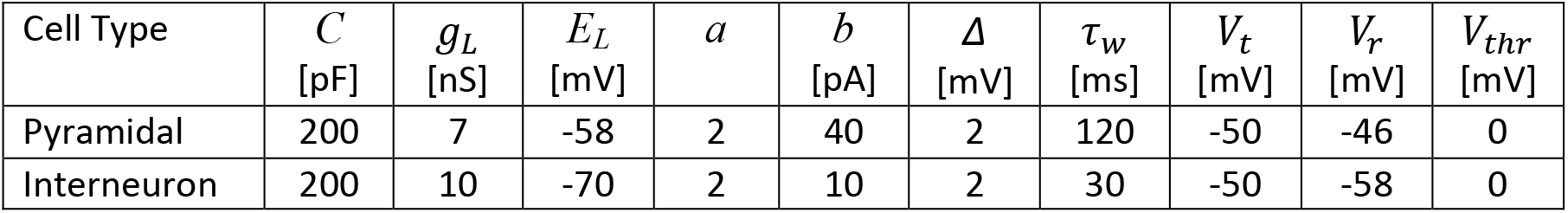
CA3 cells intrinsic parameters

#### CA3 cells input parameters

The level of noise in the network was set by a coefficient β = 80 for pyramidal cells and β = 90 for interneurons. The baseline voltage of the cells, was controlled with constant input *I_DC_*. Values of *I_DC_* were selected from Gaussian distributions with mean 24 (pA) and standard deviation 30% of the mean for pyramidal cells, and mean 130 (pA) and standard deviation 30% of the mean for interneurons.

Synaptic currents were modeled with double exponential functions, where one exponential captures the rise and the other one the decay of the synaptic conductance probability variable *s(t)*. For every cell *n* we had

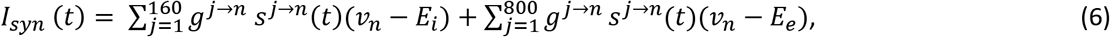

where *E_i_* and *E_e_* stand for reverse potential of inhibitory and excitatory synapses, respectively. In simulations we used *E_i_* = −80 mV and *E_e_* = 0 mV, and

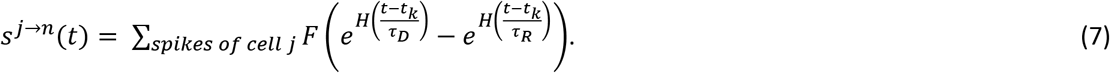

The constant *F* is a normalization coefficient, set so that every spike in the double exponential within parentheses peaks at one, and for each gate type it is a function of the time constants of rise and decay. *H*(·) represents the Heaviside function, and *t_K_* are the spike times of pre-synaptic cell *j*. The time scales of rise (*τ_R_*) and decay (*τ_D_*), expressed in ms, are given in Table 2, together with parameters controlling the strength of synapses (and hence the *g^j→n^* values), introduced in section 2.2.

**Table 2:** CA3 synaptic parameters

| Synapse Type | $\tau_R$<br>[ms] | $\tau_D$<br>[ms] | $\mu$<br>[nS] | $\sigma$<br>[nS] |
| --- | --- | --- | --- | --- |
| AMPA (Py→Py) | 0.5 | 3.5 | $34/N_{py}$ | $0.4 \mu$ |
| AMPA (Py→Int) | 0.5 | 3 | $77/N_{py}$ | $0.4 \mu$ |
| GABA<sub>A</sub> (Int→Int) | 0.3 | 2 | $54/N_{int}$ | $0.4 \mu$ |
| GABA<sub>A</sub> (Int→Py) | 0.3 | 3.5 | $55/N_{int}$ | $0.4 \mu$ |

### 2.2. Network connectivity

The model represents CA3 as a one-dimensional network. To build connections within CA3, we populated connection matrices across the different sub-population of excitatory and inhibitory neurons.

**For excitatory connections from and to pyramidal cells**, we first select a radius of about a third of all pyramidal cells in the network around the matrix diagonal (which corresponds to a radius around the pre-synaptic cell). Outside of this radius, no connections are formed. Within this radius, we shape a non-uniform probability of connection, depending on the distance between the pre-synaptic cell and the post-synaptic cell (distance in terms of their indexes in the 1D network). Specifically, we scale the decay of probability of connection with distance using a cosine function: if *x* is the index distance within the network, *y* = arctan(*kx*)/arctan(*x*) imposes the decay probability *p*(*y*) = *P* cos(4*y*), where *P* is the peak probability (*P* = 1 in our case) and *k* = 2 is a parameter controlling the decay of connection probability within the radius. As a result, the probability of connection along the diagonal is higher for cells with indexes nearby the presynaptic cell and decreases progressively with cell index distance.

**For excitatory connection from pyramidal cells to interneurons**, since the two populations have different sizes, we first identify an approximate diagonal to attribute to the connection matrix, so that every pre-synaptic pyramidal cell is assigned an inhibitory neuron spanning the whole connectivity matrix. Around that diagonal, we choose a radius about a third of the post-synaptic population (interneurons). Outside of that radius, no synapses were formed. Within that radius, we used the same distance driven scaling with the function *p*(*y*) = *P* cos(4*y*) with *P* = 1 and *y* = arctan(*kx*)/arctan(*x*), where *x* is the distance (in cell index within the interneuron population) between the post-synaptic interneuron and the diagonal interneuron point for a given pre-synaptic pyramidal cell, and *k* = 2 controls the decay connection probability within the radius of non-zero synapses.

**For inhibitory connections to both pyramidal cells and interneurons**, we identified the connection matrix diagonal and consider a radius about one third of the post-synaptic network. Within the radius, the probability of connection was uniform (*P* = 0.7) in both inhibitory projections to excitatory cells and to interneurons. Once synapses presence was established according to the aforementioned probabilities, synaptic weights for all synapse types were sampled from Gaussian distributions with variance (*σ*) given by a percent of the mean (*μ*). Parameters *μ* and *σ* used in the simulations are given in Table 2. It is to note that the mean here declared was normalized by the total number of cells in the pre-synaptic population before the variance to the mean was introduced in the distribution (*N_PY_* and *N_Int_*, respectively). Since the excitatory and inhibitory populations are of different sizes, a direct comparison of the parameter values or their magnitudes would not account for the effective values used in the simulations. One example of connection matrix for CA3 pyramidal cells is reported in Fig 1.

**Figure 1.**
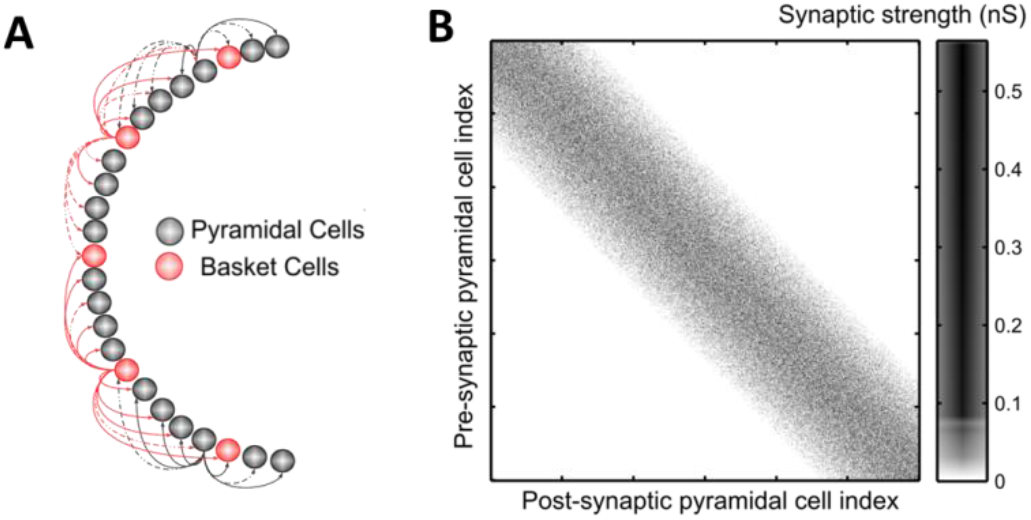
Model Connectivity. **A.** Schematic of the model of CA3, with pyramidal cells (gray) and interneurons (red), note that the network is arranged in one layer. **B.** Example of synaptic connections between CA3 pyramidal cells in one simulation. Pre-synaptic cell index is on the -axis. Note the higher density along the diagonal, decreasing toward the outermost part of the matrix. Also note that the synaptic weights (in nS) are not distributed with any topological preference.

### 2.3. Stochasticity of Network Activity

For every different simulation, we selected a new specific network connections pattern, new direct current values assigned to each cell, and delivered newly generated OU-processes traces to each neuron during each second of a simulation. To represent network activity, we studied the spike patterns in rastergrams (marking spikes by cell index in time) and the overall probability of spiking in the pyramidal cell population, which was found by finding the total spike counts in windows of 30ms length, slid every 10ms, and dividing the spike count by the size of the pyramidal cell population (1200). Whenever a sharp wave event was present in CA3, this spiking probability measure would show a rapid (highly nonlinear) rise to a peak and an abrupt decline back to baseline right after the peak. Rastergrams show how stochastic the emergence, size and location of each sharp wave can be in a simulation, and the probability of spiking traces capture the dynamics of increased firing leading up to a full blown sharp wave, which can start up to a full 150ms earlier than the sharp wave peak (Fig 2). We interpret this peak as a measure of how synchronous the sharp wave event is. The model was solved using a one-step Euler method, with integration step 0.001ms.

**Figure 2.**
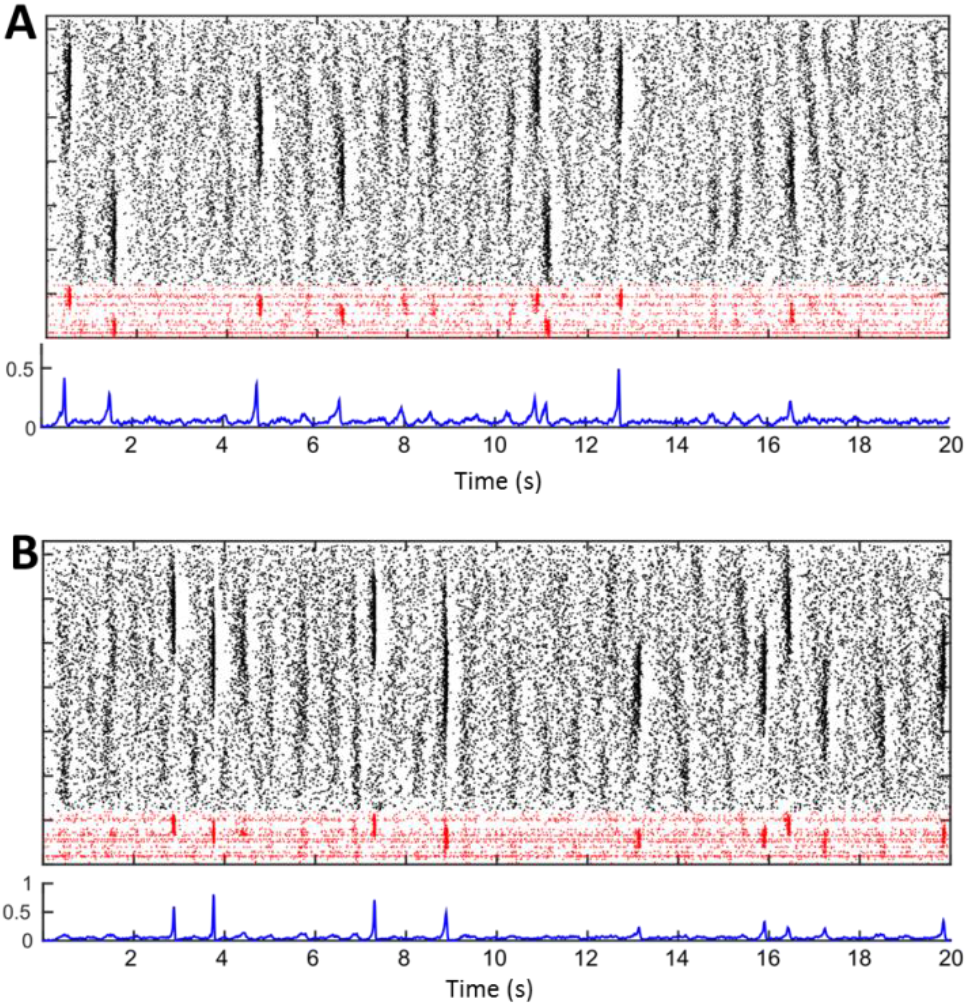
Examples of CA3 network activity in the original Model. **A.** Top plot: spiking activity in one example simulation of CA3. The rastergram shows in black dots marking spikes of pyramidal cells, and in red interneurons. On the y-axis different cells are stacked by cell index, and on the x-axis time is in seconds. The line below shows the probability of spiking for CA3 pyramidal cells in windows of 30ms. Note that high-spiking events appear as darker regions in the rastergram and as peaks of the spiking probability. The lower two plots represent a separate example of CA3 activity, generated by one new sample of connection matrix, heterogeneity DC values and ever-changing independent noise traces. Note that sharp-wave occurrence, size, and synchrony vary in time and across simulations. **B.** Same as A, but with a new instantiation of network connectivity, input noise and heterogeneity parameters

### 2.4. Instability of Network Dynamics

The rastergrams of network activity show complex dynamics of response to the input noise, including the formation and development of various sharp wave (SW) events. The input noise clearly plays a critical role in generating the complex distribution of activity patterns in the rastergrams, however the dynamical features of the network are also critical to the formation and shaping of SW. To evaluate the main contributors to the generation of the events we studied if SW were triggered exclusively by the input noisy patterns or if network dynamics also contributes the initialization of events. It is known that noise can lead to onset of synchronization that leads to event formation through the increase of reliability of spike timing of neurons [21]. One of the ways to evaluate this effect in the network is to check if the network activity in SW generation is locked to input-driven patterns. Such an analysis can be done adopting the Auxiliary System Method developed in studies of generalized synchronization of chaos [22]. This method can tell if the response dynamics of a dynamical system driven with complex signals is stable or not using an auxiliary system, which is a replica of the response system. Simulating response and the auxiliary systems, driven by the same inputs, but started with slightly different initial conditions one can detect if both systems converge to the same patterns of activity, in the case of stable response, or the initial deviations between the systems grow in time indicating the instability of the response dynamics.

The results of the auxiliary system analysis in application to the CA3 network are presented in Fig 3, where rastergrams of the network spiking of response system (the original network, Eqs. 1–5) are indicated by black dots, while spikes of the auxiliary network are shown as magenta dots on the same plot (Fig 3A). Since the timing of individual spikes is extremely sensitive to the intrinsic dynamics of neurons we do not expect onset precise synchronization between the networks at the level of individual spikes. Therefore, we focus on the analysis of spatio-temporal patterns of averaged spiking activity of the networks. We obtain such averaged activity by computing a 2-D cross-correlation of spikes both in time and in cell positions using Hanning window 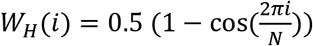, where *i* = 0,1,.*N*-1. Index *i* represents the time sample, for convolving in time with (*N*=*N_T_*=400), or cell number for computing convolution along the cell positions with (*N*=*N_P_*=80). The smoothed field of network activity patterns given by black dots is presented in Fig 3B the dynamics of perturbation computed as the difference of such fields in two identical networks shows that the errors given by absolute value of the perturbations grow in time, see Fig 3C. Note the Z-scales in the intensity plots of Fig 3B and C are the same.

**Figure 3.**
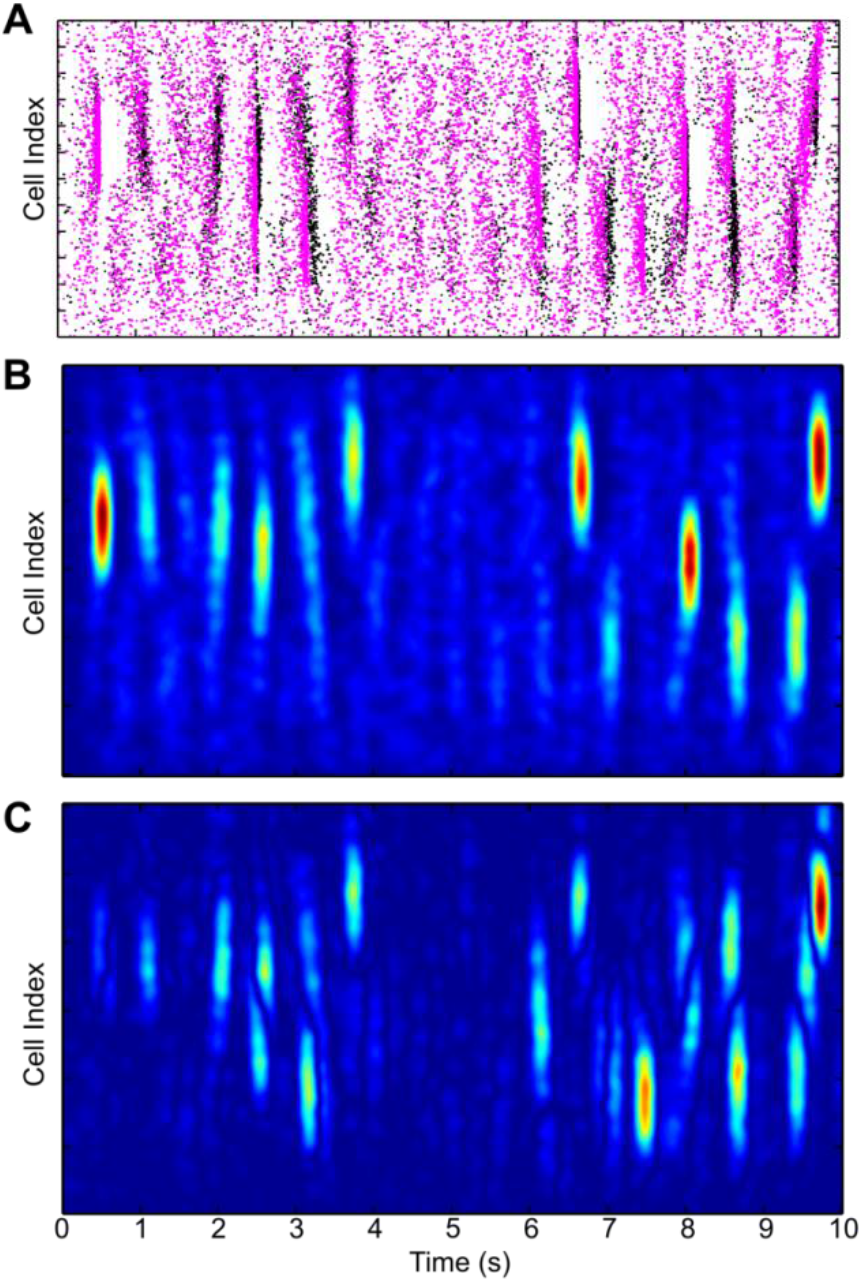
Network activity is sensitive to initial conditions. **A.** Rastergrams illustrating dynamics of PY cells spiking (black and red dots) generated by two identical networks in response to the same noisy inputs, but started at slighly different initial conditions. **B.** Smoothed actibity field of the first network shown in A with black dots. **C** Evolution of the deviations (errors) between network illustrated with the smoothed network fields demonstrate the increase of the perturbations in the network with time.

The increase of perturbations with time during a simulation indicates that network activity in the regime of generation of SW is not fully controlled by the noise inputs and that the network dynamics shows instability in its response behavior. Such instability supports variability in the network activity patterns and affects timing and shape of individual SW events. In application to our study, changing only the time step of a simulation, even if maintaining the network dynamics exactly the same, would introduce a slight perturbation of the solution at every step of integration. The instability we found in the network implies that, even in the ideal situation of same exact inputs, two identical SW networks with different integration steps could show differences in their activity responses. In the focus of our research goals it implies that criteria for matching of the SW dynamics with increasing step size should capture the capability of generation of SW events with proper characteristics of the individual events (e.g. like spiking rate profiles and fraction of neurons involved), rather than precise replication of the detailed structure of the spatio-temporal pattern generated in the response to the same sets of input patterns resampled in accordance with the step size.

## 3. Modification of the model to perform larger step simulations

To make model suitable for large step simulation we explicitly rewrite it in the form of a map using Euler 1-step integration method and changing the reset conditions for better control of spiking events. The equations will be of the form

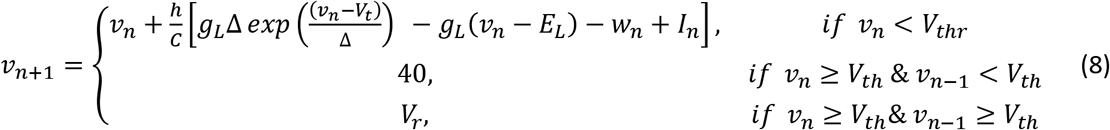

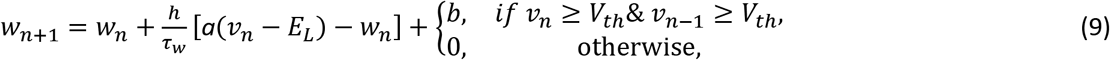

where input currents are also in discrete-time with time interval *h*:

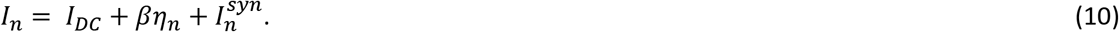

Note that the main difference between this map model and the map obtained by using fixed step Euler scheme of the original model is a modified reset condition, which now takes two steps (iterations) because of an extra condition for shaping the spike in the waveform of *v_n_*. In the Euler scheme of original model the reset to *V_r_* will take place at *v_n_* ≥ *V_thr_*. The shaping of spike by enforcing a step to value 40mV (in the new map model) helps to maintain the consistency of spiking waveform at large values of *h*. We will discuss the effect of the spike shaping in the next section.

### 3.1 Noise in discrete time model

To integrate the solution, we also introduce noise in discrete time (*η_n_*), by producing a discretized noise trace, binned with a given time step, that shows the core property of an OU-process, namely a single pole spectrum. This method follows closely strategies developed to introduce noise conductances to cell voltages *in vitro* through dynamic-clamp experiments [17, 20, 23]. To obtain such trace for a 1s time of simulation, we first construct the noise representation in the frequency domain as a complex-valued function, assigning a norm and a phase to each frequency bin. Note that the frequency axis is binned according to the frequency resolution of the trace, which depends on the duration of the noise trace, in our case 1s and the integration time step considered: df = 1/(T+dt). To each frequency bin, we assign a complex value with phase sampled normally across the unit circle and norm the value at the given frequency bin *f* of a theoretical power spectrum of an OU-process, given by 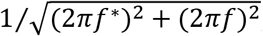; where *f\** is the location of the frequency single-pole filter, in our case 100Hz.

The complex trace on the frequency axis is then transformed to obtain a conjugate symmetric complex function, which is the representation in Fourier space of our noise sample. The inverse Fourier transform (function ifft in Matlab, The MathWorks) of this trace is the OU-sample of length 1s at the given integration time step. Since in our model we introduce a scaling parameter β which controls for the standard deviation of the noise, we further normalize the noise trace by its sample-depending standard deviation, which gives us *η_n_*.

### 3.2 Discrete time models of synaptic connection

In discrete time, the synapse rise and fall equations can be simplified using the known property that the derivative of an exponential is still an exponential. In fact, the synaptic gates (variables *s* in the equations) can be separated in two separate decay and rise terms (*p_r_* and *p_d_*), and their difference (appropriately normalized by *F*) will constitute the variable *s*. The separate terms have a straightforward ODE regulating their time evolution, which is given by 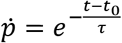, where t_0_ was the last pre-synaptic spike time; which is the same as (for *h* the integration step) 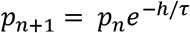. At every time step, we hence can follow the evolution of the rise and fall portion of the synaptic gate, and have *s_n+1_ = g*(*p_r,n+1_ – P_d,n+1_*). This method has been introduced in [24]. Note that this choice of discretization, while exact (which can be rarely said for discretized ODE solutions) shows explicitly the perils of using larger integrations steps: the rise and fall of a synaptic gate on different time scales could erase each other if the integration time step is too large. We keep our analysis for integration steps up to 0.5ms, which is smaller than all the decay times of synapses in the network.

### 3.3. Pyramidal cell activity when changing integration step

It is meaningful to start our evaluation of the effect of changing integration step size on cell dynamics from the spiking response to a step of current. In fact, experimental measurements that characterize a cell basic biophysical properties measure its response to step currents on f-I curves (frequency-current curves) [17]. The response of a cell to a step of current is shaped by intrinsic dynamics (for example, having a bursting response rather than a fast spiking one). It is known that the Adaptive Exponential Integrate and Fire, the type of equation our cell models are based upon, can show quite a large set of different behaviors across parameter regimes [15, 25].

Consider the effects of increasing the integration step size on the dynamics of a single CA3 pyramidal cell, activated by a rectangular pulse of injected current *I_n_*. We set the parameters of the map to be the same as in the original model with the new parameter *V_th_* = –43.5. Waveforms *v_n_* and *w_n_* illustrating a response activity evoked by a rectangular pulse of amplitude *A* = 450 started at *t_START_*=50ms and terminated at *t_END_*=250ms are presented in Fig. 4 A and B, where the waveforms were computed for original model with *h* =0.001 and map model with *h* =0.5.

**Figure 4.**
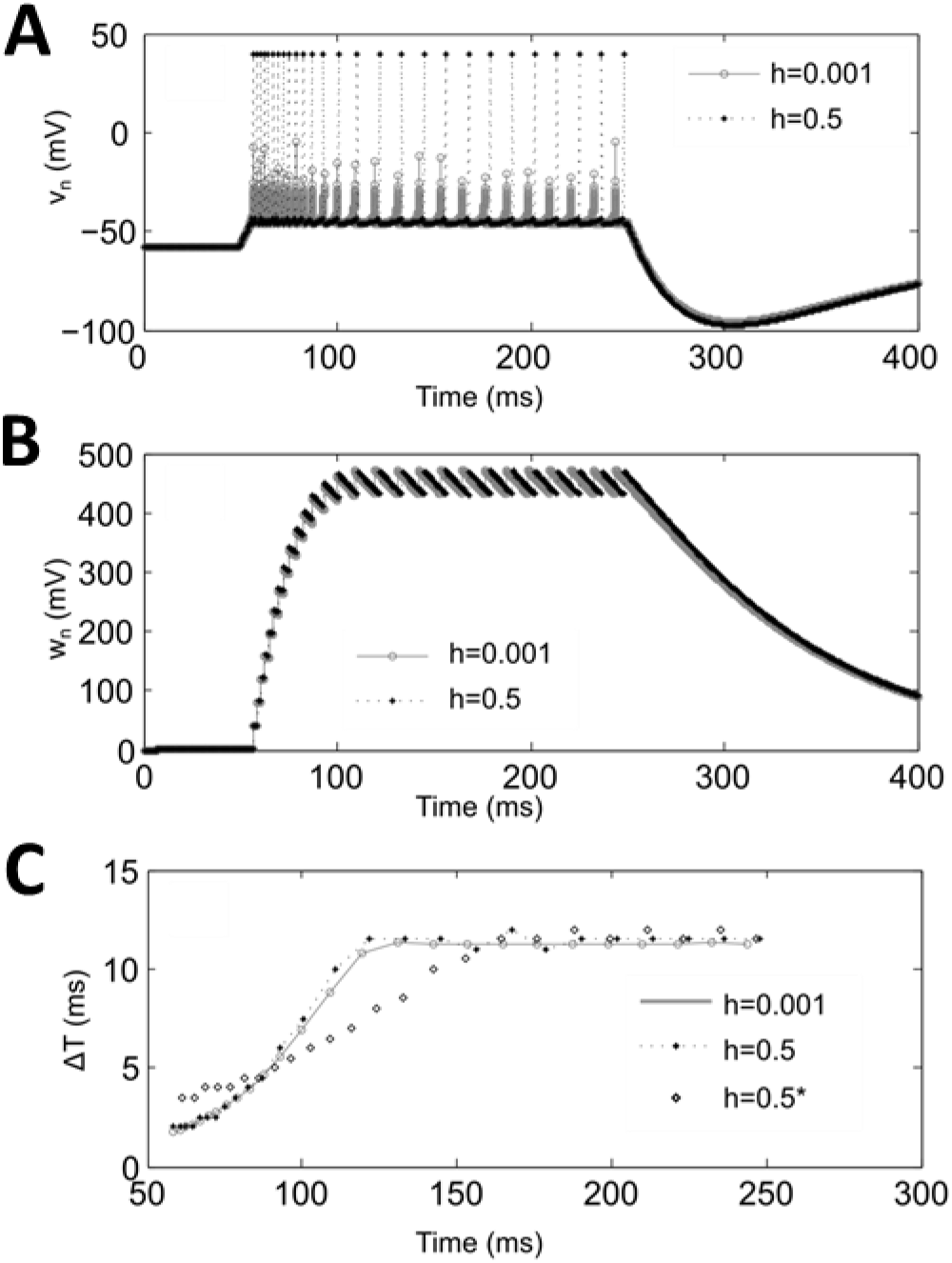
Comparison of single cell activity with different integration steps. **A-B.** Waveforms of *v_n_* and *w_n_* computed for the original model, Eqs. 1–3, with *h* = 0.001 and the map, Eqs. 8–9, with *h* = 0.5 with the same set of parameter values except for *V_th_* = –43.5. Panel **C** shows spike timing of the corresponding activity patterns plotted on the plane representing moments of spikes and proceeding inter-spike interval. Case *h*=0.5* correspondes to the spike timing obtained using the original model, Eqs. 1–3, simulated with stime step 0.5ms.

Dynamics of the spike timing in the activity patterns of each waveform is shown in Fig. 4C, which presents each spike as a point in the plane (*T_s_*, Δ*T_S_*), where *T_s_* is the time of the *s*-spike and Δ*T_S_ = T_s_ — T_s-1_*. If all spikes in of comparing activity patterns overlap, then dynamics of the patterns match. To evaluate the fitness of two comparing spike patterns *S*_1_{*T_1,1_,…, T_1,M_*} and *S*_2_{*T_2,1_,…, T_2,M_*} we design a cost function as follows. For each spike point *T_1,k_* of *S*_1_ we find the nearest neighbor spike *T_2,l_* in sequence *S*_2_ in the plane, i.e. the point that give the minimum value of the distance computed as 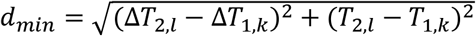. Then, for each nearest neighbor pairs we compute the mismatch between spikes as *ϵT_1,k_* = |Δ*T_2,l_* – Δ*T_1,k_*|, which characterizes error in local inter-spike intervals (ISI). We also compute the number spikes in *S*_2_, which were not selected as the nearest neighbors and normalize it to total number of spikes in *S*_2_, to get the value *ϵS_2 1_*. We repeat the same calculations with sequence of spikes *S*_2_ to get *ϵT_2,n_* and *ϵS_1,2_*. To evaluate patterns mismatch we compute total error as

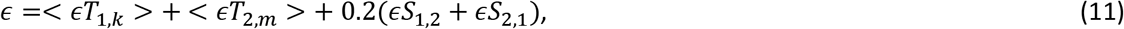

where <..> means averaging over all spikes in the sequence. Note that total error *ϵ*, representing the mismatch between two patterns, in our case does not include mismatch in simultaneous firing of the comparing nearest neighbor spikes directly. This allows us to deal with the cases of tonic spiking settled in the case of depolarizing DC input, when spikes in compared models can have some phase shift despite the precise matching of the spiking rate. The coefficient 0.2 in equation for *ϵ* is a weight, which is selected empirically to get a better consistency in changes of cost function shape, *ϵ* vs a parameter of the map.

Using *ϵ* as a measure of mismatch between comparing patterns of spiking activity in the models evoked by the same input, we consider how accuracy of the original model evolve as we change the size of *h* focusing on the analysis of cost function *ϵ(V_thr_)*. The results of such analysis is shown in Fig 5A. One can see that even a small change of *h* (e.g. 10%) significantly increases the value of errors *ϵ* in the original model, where *V_thr_* = 0. The presented plots also show that if the value of *h* increases beyond 10%, than for a given value of *h* one can find an optimal value of *V_thr_*, which minimizes the deviation of spiking patterns from the original model, Eqs.1–3, computed with *h* = 0.001 and *V_thr_* = 0. Fig 5A also shows that the use of additional spike shaping iteration, Eqs. 8–9, affects the optimal values of *V_thr_*. Compare plots for *h* = 0.005 and *h* = 0.005(*s*), where *(s)* indicates the use of spike shaping condition.

**Figure 5.**
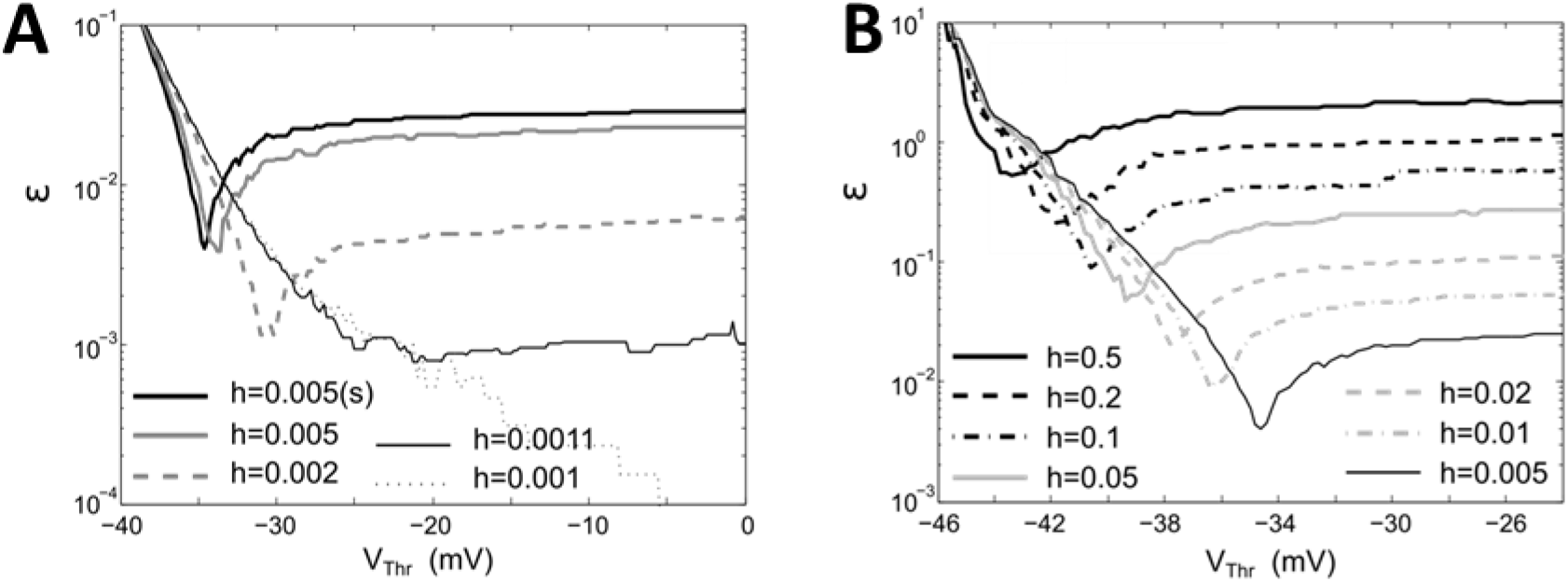
Shapes of cost function Eq. 11 computed for different values of *h*. **A.** Original model, Eqs. 1–3, with rectangular pulse size *A* = 450 integrated with fixed step Euler scheme **B.** Modified map, Eqs. 8–9, simulated with *A* = 450. For comparison panel **A** includes a plot computed for modified map with *h* = 0.005 and *A* = 450 labled as *h* = 0.005(s).

A typical change of cost function in modified map with increasing value of *h* is illustrated in Fig. 5B, where the plots of *ϵ(V_th_)* were computed for the spike patterns evoked by rectangular pulse of amplitude *A* = 450 and the other parameters are the same as in Fig.4. The plots show that errors grow and optimal value of *V_th_* moves down as the values of *h* increases.

### 3.4. CA3 network activity when changing integration step

To compare the simulation outcomes when changing integration step, we kept unchanged the specific network connection matrix and assignment of direct current value to each cell in the network. For the input noise received by each cell, we froze each trace sampled at the highest rate (h = 0.001) and downsample the trace to the new integration step. Within these constrains, we run copies of the “same” simulation for integration steps progressively increasing (Fig 6).

**Figure 6.**
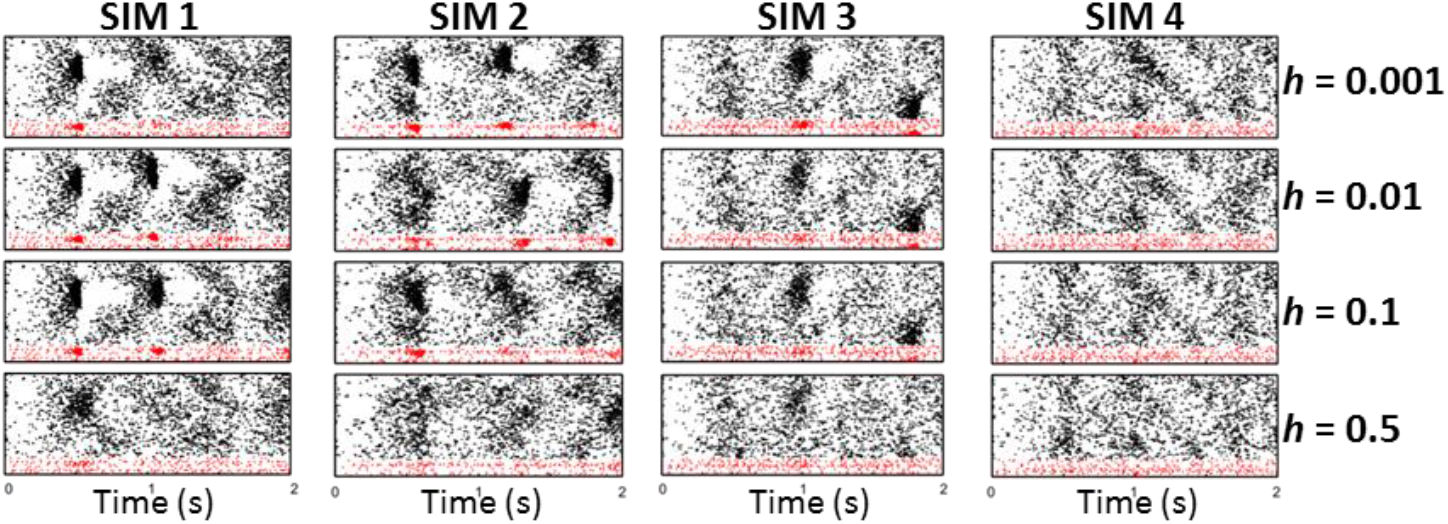
Integration step affects network dynamics. Increasing the integration step (*h*) leads to loss of sharp wave activity in the network: four examples. Across the different integration steps, all connections and heterogeneity DC values were kept identical, to test whether new integration steps would lead to different overall network patterns.

In Fig 6, examples show how for a network of CA3 pyramidal and basket cells, increasing step size affects the presence of spontaneous sharp wave activity. To test specifically what changed in network dynamics, we created an initial pool of ten simulations, each 10s long, for the finest integration step (*h* = 0.001 ms). We run a copy of each simulation in this group where we only changed the integration step to h = 0.5ms. We then compared a number of biologically relevant characteristics of the SW events in the original model (*h*=0.001ms) and in the large integration step case (*h*=0.5). In each simulation, we counted the number of SW detected (SW count), the duration of times in between two successive SW (inter-SW times), the fraction of pyramidal cell population which spiked at least once in each SW (SW size), the number of spikes emitted by cells during a SW (spike count in SW), the peak value reached during a SW of the 30ms-windowed probability of spiking for pyramidal cells (SW synchrony, i.e. the peak values of the blue lines in Fig 2) and the duration of SW events (SW duration). Across all simulations, we pooled these values to create histograms of the statistics (Fig 7). It is to note that the statistical distributions of these properties were carefully introduced in the original model to resemble known biologically quantified SW properties [26–28], so it is important that they are preserved in solutions for larger integration steps.

**Figure 7.**
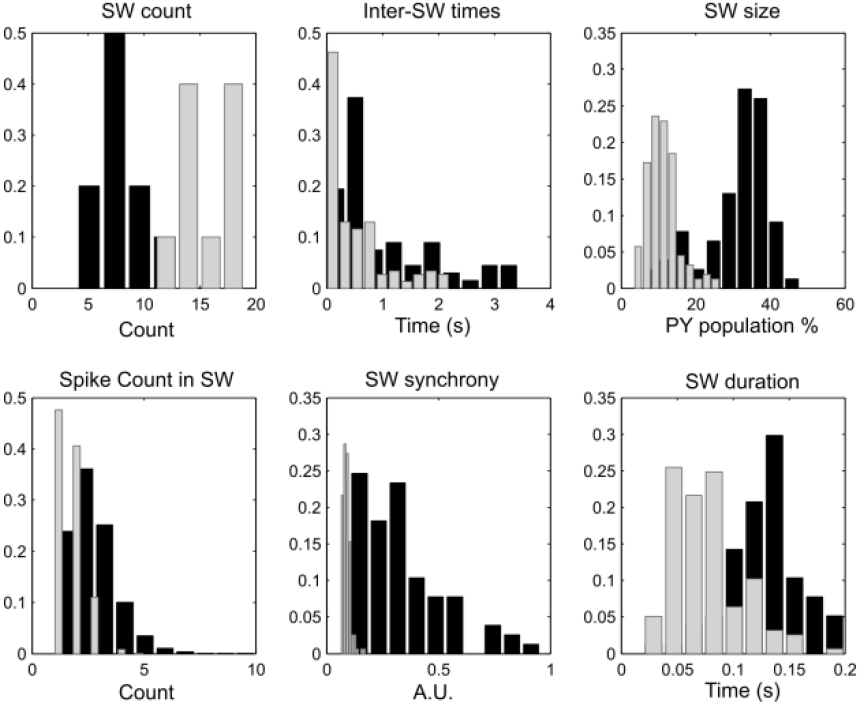
Integration step affects the statistics of SW properties. Panels show histograms of SW properties with data collected across 10 simulations, each 10s long. Histograms of the statistics for the small integration step (*h* = 0.001 ms, black bars) and for the large integration step (*h* = 0.5 ms, gray bars). SW count: count of SW in each simulation. Inter-SW times: time in between the end of a SW and the beginning of the next one. SW size: fraction of the pyramidal cell population which spikes at least once in the SW. Spike Count in SW: number of spikes released by the pyramidal cells active during SW. SW synchrony: peak height of the probability of spiking of the pyramidal cell population (in 30ms time bins). SW duration: time between start and end of each SW.

As can be seen in Fig 7, increasing the size of integration step led to too many SWs. This might seem to contrast with our observation in Fig 6, where we show that increasing the integration step led to loss of SW. In fact, this apparent contradiction can be reconciled by a small technical point. The automatic detection of SW events is based on the time evolution of the network-wide probability of spiking of pyramidal cells (blue lines in Fig 2), binned in 30ms windows and sampled every 10ms (*p_E_(t)*), which we use to compute the SW synchrony, given by the peak value of this probability during a SW. If a network activity profile with no sharply defined SW event is passed through the algorithm, the “ghosts” of SW activity might lead the probability measure to occasionally cross the detection threshold, which is given by *p_Th_ = b_E_ +2σ*, where *b_E_* is the baseline of pyramidal cell activity, found as the mean value of all *p_E_(t)* values that are below the sum of mean and standard deviation of *p_E_(t)*. As a consequence, the baseline is close to the mean of *p_E_(t)* but always below it. The threshold *p_Th_* is reached when crossing the sum of baseline and σ, which is the standard deviation of *p_E_(t)*. Hence, every simulation has its own baseline and threshold, and simulations without any SW synchronous enough to drive the creation of a meaningful threshold can detect as SW events which are not synchronous or large. This is exactly the case with the *h*=0.5ms simulations, as can be verified in the “SW synchrony” panel of Fig 7, showing all the SW detected in the large integration step case (in red) tightly packed near zero synchrony.

Other discrepancies in the *h*=0.5 simulations are given by SW which are too small (SW size panel) and last about half the appropriate duration (SW duration panel). Overall, increasing the time step without any form of adjustment led to a dramatic loss of SW-like behavior in the model activity. We believe that this is due to the loss of excitatory propagation in the network, because increasing the step size hyper-synchronizes some sparse spikes, which cannot accumulate in time on their shared targets, therefore losing the slowly building excitatory activity which precedes every SPW by about 50-100ms (biologically consistent).

Since larger integration steps caused the CA3 network activity to lose (for the most part) the ability to start sharp waves, even when preserving all other network properties as connectivity, initial conditions and frozen noise, we chose to adjust some parameters to test whether we could recover the SPW spontaneous emergence. Since we thought that excitatory synaptic transmission was affected in the larger integration step cases, we investigated the effects of strengthening two network parameters which we knew enhanced the likelihood of SW activity in the network when simulated at a fine time step: the strengths of excitatory synapses and the standard deviation of the noise delivered to excitatory cells (both mechanisms efficient at promoting spiking in the CA3 population).

Specifically, in each of the original 10 simulations (blue histograms in Fig 5) we multiplied all the pyramidal-to-pyramidal synaptic connections by a non-dimensional scaling factor *G_α_* which varied between 1 and 1.04. Also, we multiplied the factor β which scales the contribution of the noise current to pyramidal cells (see the system of equations introduced in section Cells: equations and parameters) by a factor *B_α_* which varied between 1 and 1.05. As a result, these values were either unaltered (at scaling 1) or slightly increased. The two scaling operations were carried out independently, generating a two-dimensional parameter space in which, for every combination of *G_α_* and *B_α_*, we run the 10 simulations with integration step *h*=0.5ms.

We pooled data from simulations to compare SW properties in the original model solved with small integration step and the modified models solved with large integration step. We used the same properties that we introduced in Fig 7: SW count, size, duration, synchrony, the Inter-SW times and the cell spike count during SW events. When comparing across all possible scaling combinations, we found one which performed the best in terms of matching the distributions of these properties in the pool of simulations. It was given by *G_α_* = 1.025 and *B_α_* = 1.04, and the resulting statistics are shown in Fig 8. In Fig 9 we show one example of the rastergrams for the same simulation in the original model, increased integration step and modified model conditions.

**Figure 8.**
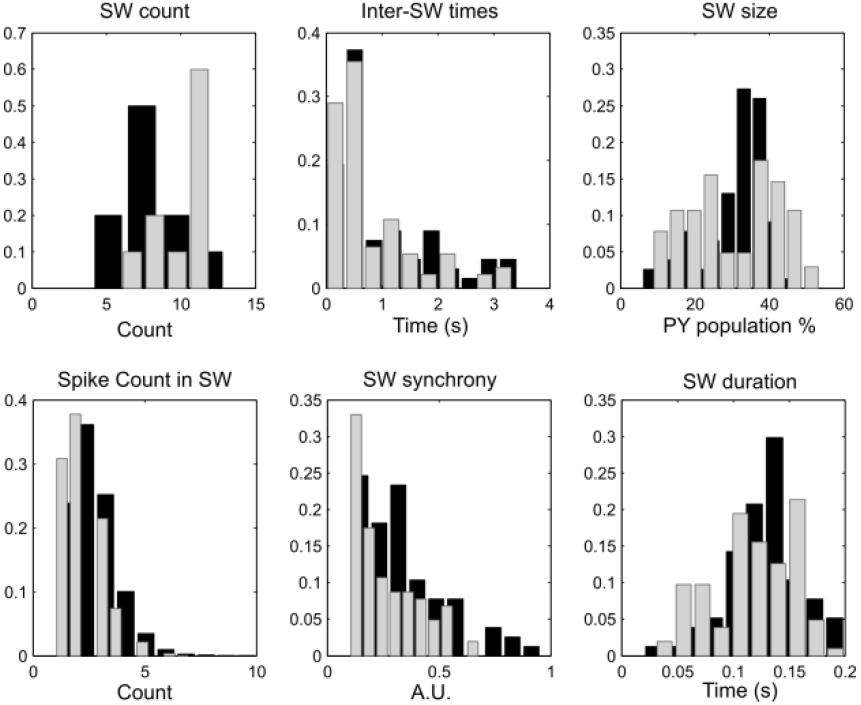
Adjustments to simulations parameters rescue the SW dynamics with large integration step. In the panels, computed as in Figure 7, we should a comparison between data pooled by simulation of the original model (black bars) solved with *h*= 0.001ms and the modified model (gray bars) solved with *h*=0.5ms. Note that while not perfectly overlapping, all distributions lie in overlapping ranges and show similar trends.

**Figure 9.**
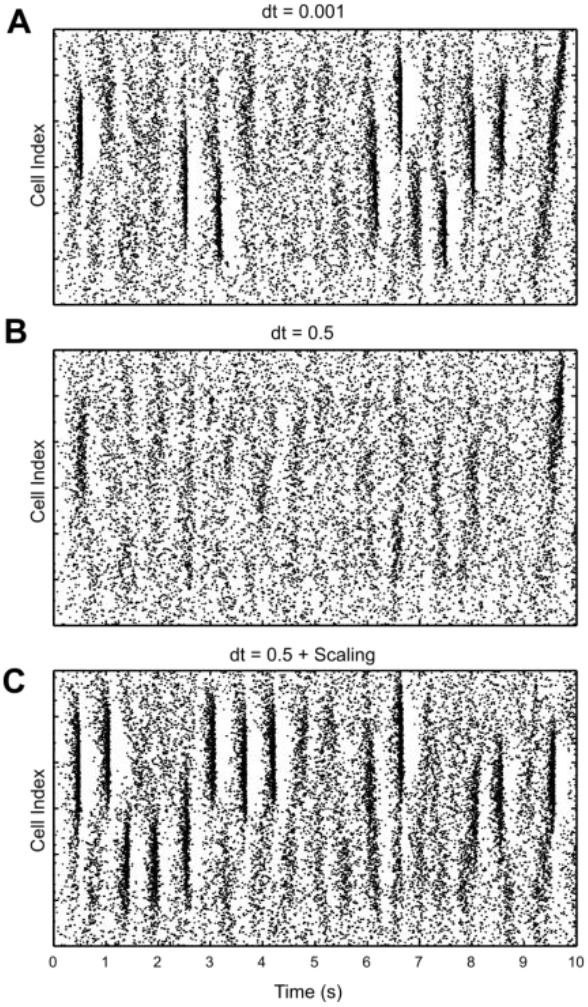
Example of recovered network activity at large *h*. Shown is one simulation solved at *h*=0.001ms (**A**), the same network at *h*=0.5ms without any other changes (**B**). **C**. The same network (*h*=0.5) with re-scaled excitatory connections and excitatory noise size.

## 4. Conclusions

We have studied how an ODE-based phenomenological model of spiking neuron, known as Adaptive Exponential Integrate and Fire Model, can be modified into a form of discrete-time model (a map) that tolerates simulations with large time step and supports proper behavior of network dynamics in generation of sharp-wave oscillations modeling CA3 region of hippocampal network. We have demonstrated that dynamics of oscillations produced by considered network models is very sensitive to the variation of parameters of the neurons and synapses forming the network, changes in the noise and time step of simulations used in Euler scheme of integration. To achieve reliable operation of network in the regime of sharp-wave generation the integration step should be set at very small values ∼0.001ms, which makes difficult to apply this model for detailed analysis of large-scale network dynamics.

Since noise plays a critical role in the initiation of sharp waves in this model, it has to be properly modified to support proper network activity as the integration step changes. We have shown that even for the same noisy inputs (frozen noise) used in the simulations the response behavior of the network model is very sensitive to small variations in initial conditions indicating that although the network reacts to various noisy events in inputs whose coherency can evoke a sharp wave, the network dynamics has intrinsic instability. This instability hampers the tuning of the model to achieve behavior matching between models simulated with different step size h. Therefore, tuning of the model with large time step is done by matching the spiking response activity in the individual neurons. We have shown the most appropriate parameter to adjust here is the value of threshold level. A similar approach has been also successfully used recently for control of sampling in a map-based model of spiking neuron [29].

Additional parameter turning with increase of time step has to be done at the level of network simulation. This is mostly needed to properly balance the effects of noise and synaptic inputs on the network responsivity. We used strengths of the synaptic connections as control parameters of the network tuning. Because of intrinsic instability of the network dynamics capable generation of sharp-waves the only working indicators of similarity between the networks, we found is the statistical analysis of SW events. This analysis includes distributions of: SW count, Inter-SW times, SW size, Spike count in SW, SW synchronization and SW durations, which was kept within biological constrains [28].

## Acknowledgements

This work was supported by grants from ONR (MURI: N000141310672).

## Footnotes

1 Abbreviations CA3, CA1: Cornus Ammonis area (1 and 3, respectively). Anatomic sub-regions of the hippocampus. SWR: sharp-wave ripples. Hippocampal events in area CA3 and CA1, tied to sleep-dependent memory consolidation.

